# Investigating RNA splicing as a source of cellular diversity using a binomial mixture model

**DOI:** 10.1101/2023.10.17.562774

**Authors:** Keren Isaev, David A Knowles

## Abstract

Alternative splicing (AS) contributes significantly to RNA and protein variability yet its role in defining cellular diversity is not fully understood. While Smart-seq2 offers enhanced coverage across transcripts compared to 10X single cell RNA-sequencing (scRNA-seq), current computational methods often miss the full complexity of AS. Most approaches for single cell based differential splicing analysis focus on simple AS events such as exon skipping, and rely on predefined cell type labels or low-dimensional gene expression representations. This limits their ability to detect more complex AS events and makes them dependent on prior knowledge of cell classifications. Here, we present Leaflet, a splice junction centric approach inspired by Leafcutter, our tool for quantifying RNA splicing variation with bulk RNA-seq. Leaflet is a probabilistic mixture model designed to infer AS-driven cell states without the need for cell type labels. We detail Leaflet’s generative model, inference methodology, and its efficiency in detecting differentially spliced junctions. By applying Leaflet to the Tabula Muris brain cell dataset, we highlight cell-state specific splicing patterns, offering a deeper insight into cellular diversity beyond that captured by gene expression alone.

## 1 Introduction

Alternative splicing (AS) is a crucial cellular process that generates multiple RNA isoforms, leading to the protein and regulatory RNA diversity necessary for the proper function of complex organisms [1, 2]. The mammalian nervous system is a prime example where AS is prevalent and responsible for increased proteomic and functional complexity [3, 4, 5]. However, the extent to which AS contributes to cell-type level specific variation remains incompletely understood. Addressing this requires an unbiased analysis of AS in relation to cellular states yet quantifying it using common single-cell RNA sequencing techniques such as 10X is challenging due to its pronounced 3’ bias in coverage. Smart-seq2 however, while less high-throughput than droplet based approaches, offers superior transcript coverage, making it a promising experimental approach [6]. Importantly, several single-cell atlases encompassing both 10X and Smart-seq2 data across various cell types and tissues in mouse and human have been published [7, 8]. These offer potential insights into the AS profiles of individual cells and cell types.

Despite the challenges posed by current single-cell technologies, a number of methods have been developed to capture differential splicing in single cells. BRIE2 [9] uses Bayesian regression to model the association of specific AS events with cell-level phenotypes. However, it is mainly tailored for exon-skipping events and requires predefined binary cell type labels such as treatment status. Another approach, scQuint [10], integrates a method where introns sharing 3’ splice sites converge to form alternative intron groups. It then maps the genome-wide intron usage onto a low-dimensional latent space with a Dirichlet-Multinomial VAE, followed by a differential splicing test across predefined cell types or clusters, which emerge from the splicing latent space. However, the obtained splicing latent space with this approach did not reveal any novel cell sub population in Tabula Muris. Meanwhile, Psix[11] employs a probabilistic model to detect splicing variation across a spectrum of single cells. Inspired by autocorrelation, Psix seeks patterns of AS that correspond to continuous cell states, such as developmental stages, without the need for explicit cell clustering, labeling, or trajectory inference. However, Psix’s approach relies heavily on a gene expression-based low-dimensional representation. Thus, it is inherently limited to identifying splicing events that are directly associated with gene expression changes. Furthermore, a significant limitation of Psix is its reliance on K-nearest neighbors (KNN) for determining cell similarity, which makes the method time-consuming, especially with large single cell datasets. Overall, the focus of recent approaches has largely remained on identifying variable cassette exons, likely resulting in the oversight of more complex splicing events.

To address this gap in our understanding of splicing events in single cells, we developed Leaflet, building on the splicing quantification representation employed in Leafcutter [12] for bulk RNA-seq. Our binomial mixture model uses junction counts from split reads (described in **Section 2)** to model latent cell states and the splicing patterns that define them **(Figure 1)**.

**Figure 1:**
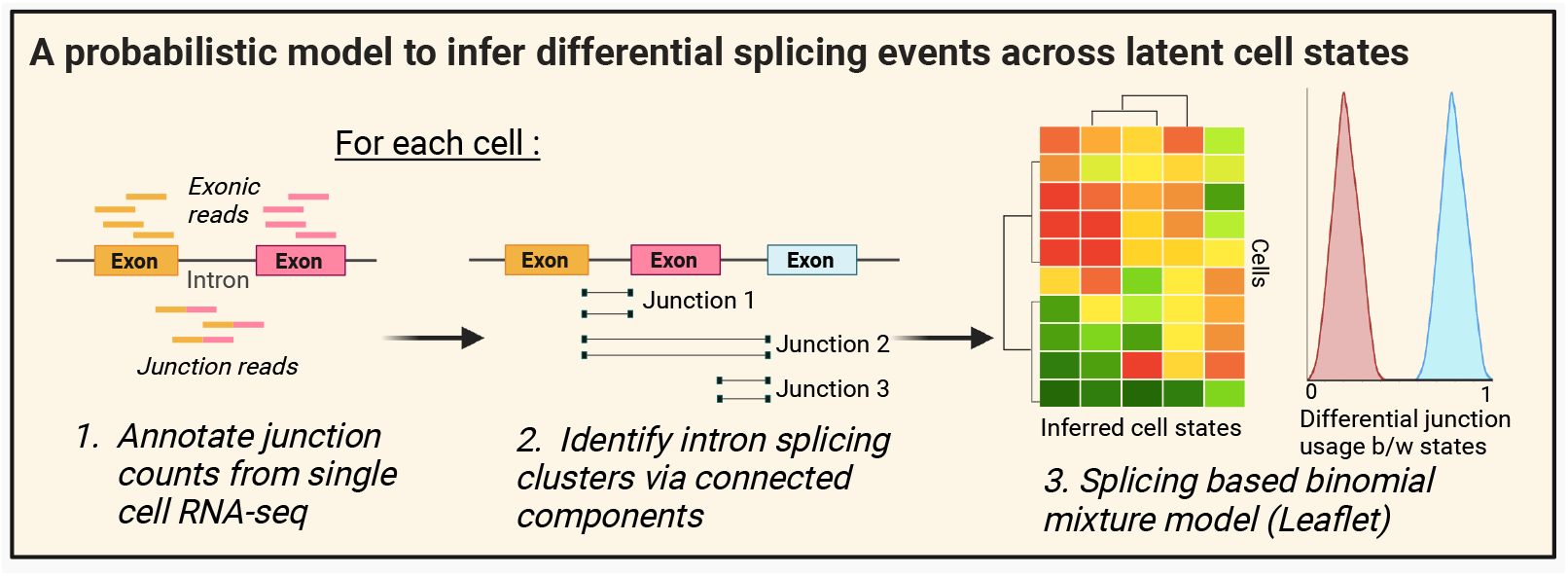
Overview of single cell AS quantification. Junction coordinates and reads are first obtained for each cell, followed by intron clustering of overlapping junctions with common splice sites. Three exemplary junctions are shown for one intron cluster event, exon skipping. Read counts for junctions and intron clusters are used as input for Leaflet’s binomial mixture model.

## 2 Splicing quantification with Smart-seq2 reads

To generate an input dataset for our model, we processed reads mapping to splice junctions in individual cells. We used Regtools [13] to obtain a matrix of junction coordinates and read counts while keeping track of cell barcodes. Leaflet follows Leafcutter’s approach to defining local splicing variation via “intron clusters” but provides additional functionality for single-cell applications. For example, in our analysis of Tabula Muris, we required each junction to be observed in at least five cells in each cell type with at least one read each. These parameters are user adjustable and the full list of them is described in the package https://github.com/daklab/Leaflet. Leaflet then combines junctions into intron clusters based on overlapping introns and splice sites. An example of an intron cluster composed of three overlapping junctions is shown in **Figure 1**. Intron clusters with a minimum of two junctions are kept for downstream analysis. We define the junction usage ratio (or in a slight abuse of notation, the percent spliced in or PSI) as the splice read count for the junction divided by the total count for the cluster.

## 3 Leaflet probabilistic model

### 3.1 Generative model

The graphical model for Leaflet is shown in **(Figure 2)** with *C* cells, *J* junctions and *K* states. Leaflet posits the existence of *K* cell states, each defined by a vector of junction usage ratios *ψ* _*k*:_. In the context of a collection of single cells, we model the prior proportion of each cell type or cell state in the sample as *θ*. Cell assignments *z*_*c*_ are sampled from a categorical distribution parameterized by *θ. z*_*c*_ along-side *ψ*_*kj*_ model the observed junction counts *y*_*cj*_. *T*_*cj*_ are the observed intron cluster counts for a given junction *j* in a given cell *c*.

**Figure 2:**
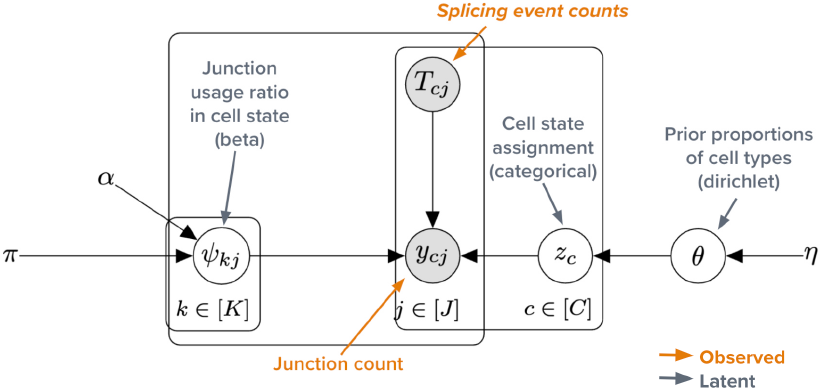
A graphical model for Leaflet’s mixture model. Variables described in **Table 1**.

Our likelihood model is

**Table 1:** Summary of Variables in Leaflet.

| Variable | Description |
| --- | --- |
| $C$ | Total number of cells in the dataset. |
| $J$ | Total number of junctions considered. |
| $K$ | Total number of cell states or cell types. |
| $\psi_{k:}$ | Vector of junction usage ratios for each state $k$. |
| $\theta$ | Proportion of each cell type or cell state. |
| $z_c$ | Cell assignment for cell $c$. |
| $y_{cj}$ | Observed junction counts for junction $j$ in cell $c$. |
| $T_{cj}$ | Observed intron cluster counts for junction $j$ in cell $c$. |

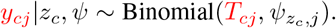

The proportion of counts attributed to each junction within an intron cluster is specified by *ψ*_*kj*_, the latent success rate of junction *j* in whichever state *k* the cell is in. The junction usage ratios are given a Beta prior distribution with fixed hyperparameters *a* and *b*,

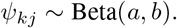

The latent cell assignments *z*_*c*_, specifying the state of cell *c*, are modeled as a categorical given *θ*, Categorical(*θ*).

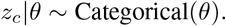

Of note, within a single cell *c*, the model assumes that the observations at different junctions are conditionally independent, given the cluster assignment and the associated *ψ*. Meanwhile, *θ*, denotes the prior on cell assignments and is modeled using a Dirichlet distribution with fixed prior *η/k*,

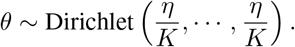

The true number of cell states is typically unknown. Our approach is to set the parameter vector of the Dirichlet to [*η/K*,, *η/K*] which approximates a Dirichlet Process (DP) as *K · → ∞*. When used for clustering, the approximate DP with large *K* provides a way to infer the number of clusters from the data, instead of having to pre-specify it. Thus, we recommend setting *K* to a larger number of cell states than expected to infer the optimal value.

## 4 Variational inference

We approximate the posterior distribution using mean-field coordinate ascent variational inference (CAVI) [14]. To learn the parameters, we optimize the Evidence Lower Bound (ELBO, 10.1). Our variational posterior is

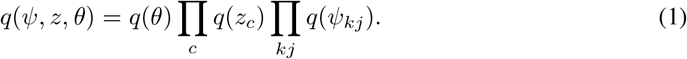

where

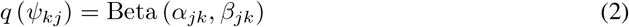

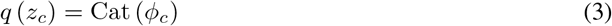

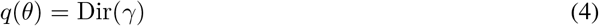

Here *α ∈* ℝ^*K×J*^, *β ∈* ℝ^*K×J*^, *ϕ ∈* ℝ^*K×N*^, and *γ∈* ℝ^*K*^ are the variational parameters to be optimized. Given the variational parameters *α* and *β*, the expected log-likelihood under the variational approximation of observing *y*_*ck*_ under component *k* is

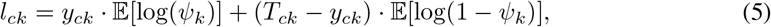

where 𝔼 [log(*ψ*_*k*_)] = digamma(*α*_*k*_) ™digamma(*α*_*k*_ + *β*_*k*_) and 𝔼 [log(1 − *ψ*_*k*_)] = digamma(*β*_*k*_) − digamma(*α*_*k*_ + *β*_*k*_).

The CAVI updates are

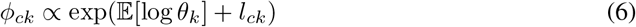

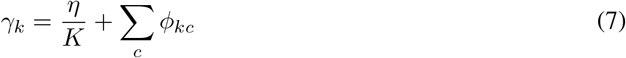

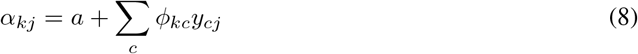

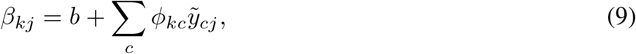

where 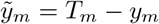 and 𝔼 [log *θ*_*k*_] = digamma(*γ*_*k*_) − digamma(Σ_*k*′_ *γ*_*k*′_). The total cluster count *T*_*cj*_ is 0 for most (cell-junction) pairs across any (current) single cell sequencing platform. It is thus highly beneficial to represent *y* and 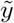 as sparse matrices. The updates for *α* and *β* are then given in matrix notation as,

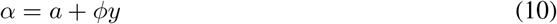

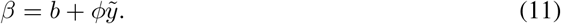

Our implementation uses `pytorch` allowing the CAVI updates to be easily run on a GPU for highly efficient model fitting.

## 5 Leaflet differential splicing analysis

We test whether junction *j* is differentially spliced across cell states by calculating an approximate log Bayes factor (ALBF) between hypothesis *H*_0_ that *ψ*_*jk*_ = *ψ*_*j*_ for all *k* and *H*_1_ that the *ψ*_*jk*_ are different across states. Given that the variational posterior on *ψ* is a Beta distribution, we can compute an approximate marginal likelihood under *H*_0_ as

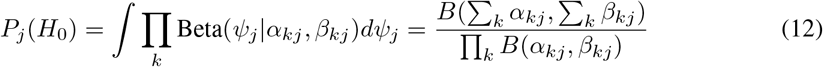

where B(*a, b*) = Γ(*a*)Γ(*b*)*/*Γ(*a* + *b*). The approximate marginal likelihood for *H*_1_ is

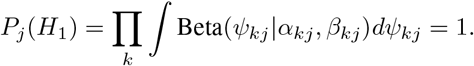

Thus, the ALBF for junction *j* is

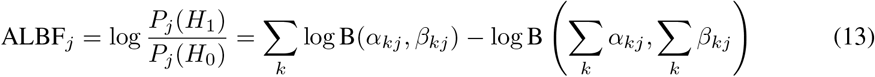

## 6 Evaluation of performance on simulated data

We compared Leaflet, Psix and Kruskal-Wallis in their abilities to detect simulated differentially spliced events. Here we show results simulating discrete cell-types, but we also compare the methods on a continuous trajectory dataset simulated by Psix in Appendix 10.2, Table 3.

**Table 2:**
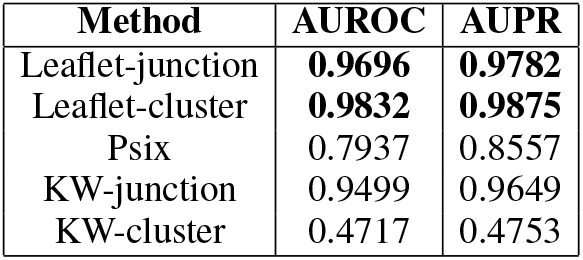
Comparison of methods on discrete counts generated from Leaflet generative model simulation. Data corresponds to 5,968 cells and 5,676 junctions from 1,892 intron clusters.

| Method | AUROC | AUPR |
| --- | --- | --- |
| Leaflet-junction | **0.9696** | **0.9782** |
| Leaflet-cluster | **0.9832** | **0.9875** |
| Psix | 0.7937 | 0.8557 |
| KW-junction | 0.9499 | 0.9649 |
| KW-cluster | 0.4717 | 0.4753 |

**Table 3:**
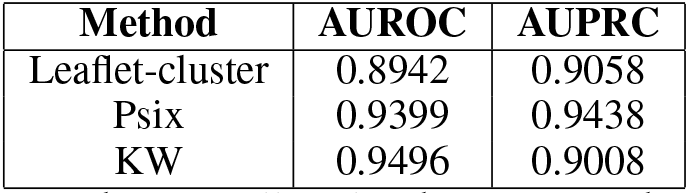
Performance at detecting differentially spliced genes from data simulated by Psix with cells along a continuous trajectory. KW: Kruskal–Wallis test.

### 6.1 Splicing differences between discrete cell-types

Our simulation approach adheres closely to Leaflet’s generative model (**Figure 2**), but uses the observed cluster counts matrix (detailed in **Section 2**) and predefined brain cell types from Tabula Muris to ensure the overall sparsity of read coverage is maintained. To allow for a fair comparison with the type of splicing events captured by Psix (exon skipping), we filtered the Leaflet intron clusters to exclusively encompass exon inclusion/exclusion events, corresponding to three junctions per intron cluster.

In our simulation, every intron cluster is randomly assigned a positive (has differential splicing across cell states) or negative label (no differential splicing). Junctions are sorted into the two corresponding to exon inclusion (**J1** and **J2**) and the one corresponding to exon exclusion (**J3**). For negative intron clusters we sample one shared *ψ*^*J*3^ from Beta(0.5, 0.5) (hyperparameters chosen to emulate the known bimodality of splicing patterns [15]) for all cell states *k*. For positive intron clusters, we sample a different 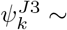 for each cell state *k*. We then set 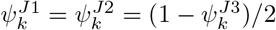for each *k*. Junction counts are then sampled from binomial distributions with these *ψ* values and total count from the observed (“real”) Tabula Muris data.

### 6.2 Identifying cell-state associated splicing events

For our first test, we focused on the two most prevalent cell types in Tabula Muris brain tissue: microglia and oligodendrocytes (5,499 cells total). From 1,892 intron clusters, we sampled split read counts for 5,676 junctions based on these specified cell types. The ability of Leaflet, Psix, and the Kruskal-Wallis test to detect the simulated differentially spliced events was then assessed.

Leaflet’s mixture model was fit on the simulated counts with 100 CAVI iterations across 50 independent trials with *K* set to two. These trials represent random initializations of the variational parameters. The trial resulting in the highest ELBO was used for downstream analysis. For each junction cell-type pair, the posterior mean was 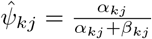. The Spearman correlation between the inferred and simulated Δ*ψ* between the two cell states was 0.92.

For comparison, Psix (version 0.11.0) was executed using its default parameters, and the first two principal components were determined based on the simulated intron cluster counts. In this setting, an intron cluster corresponded to a “gene”. Psix requires several input files including a gene expression-based low-dimensional representation, a junction counts matrix and cassette exon annotations to compute Psix scores. Empirical *p*-value estimation with its kNN-based method extends its run time (21.34 seconds for Leaflet vs 5 minutes for Psix for the same dataset).

The Kruskal-Wallis (KW) test was run using the two vectors of simulated junction usage ratios (KW-junction) or intron cluster based PSI corresponding to *J*_3_ (KW-cluster) only for the two cell types. The results are shown in **Table 2** where Leaflet consistently outperforms other methods. Specifically, Leaflet-cluster, which evaluates the overall intron cluster Leaflet score (by summing scores across the three junctions in a given cluster), achieves the highest Area Under the Receiver Operating Characteristic (AUROC) and Area Under the Precision-Recall (AUPR) scores.

### 6.3 Inference of cell state identity

Leaflet simultaneously infers junctions that are differentially spliced across cell states as well as *z*_*c*_, the cell assignments. We evaluated Leaflet on a larger simulated dataset using eight brain cell types from Tabula Muris, fixing cell intron cluster counts and true cell-type assignments while simulating junction *ψ*.

This data consists of 7,816 cells and 3,613 intron clusters corresponding to cassette exon events. We applied Leaflet’s mixture model to the simulated counts, running 100 CAVI iterations from 50 random initializations with *K* set at 50. The variational parameter *ϕ*_*c*_ determined the cell state assignments for each individual cell. To evaluate reproducibility, we analyzed the consistency of cell-cell co-assignments across random initializations **Figure 3**. This revealed a robust structure: cells originating from the same cell types were largely co-assigned across trials (Adjusted Rand Score = 1.0). Further, even with *K* set at 50, Leaflet identified the correct number of predefined cell types as eight, specifically where *θ ≥* 0.05. This analysis reflects Leaflet’s ability to detect inherent cell types by leveraging AS patterns.

**Figure 3:**
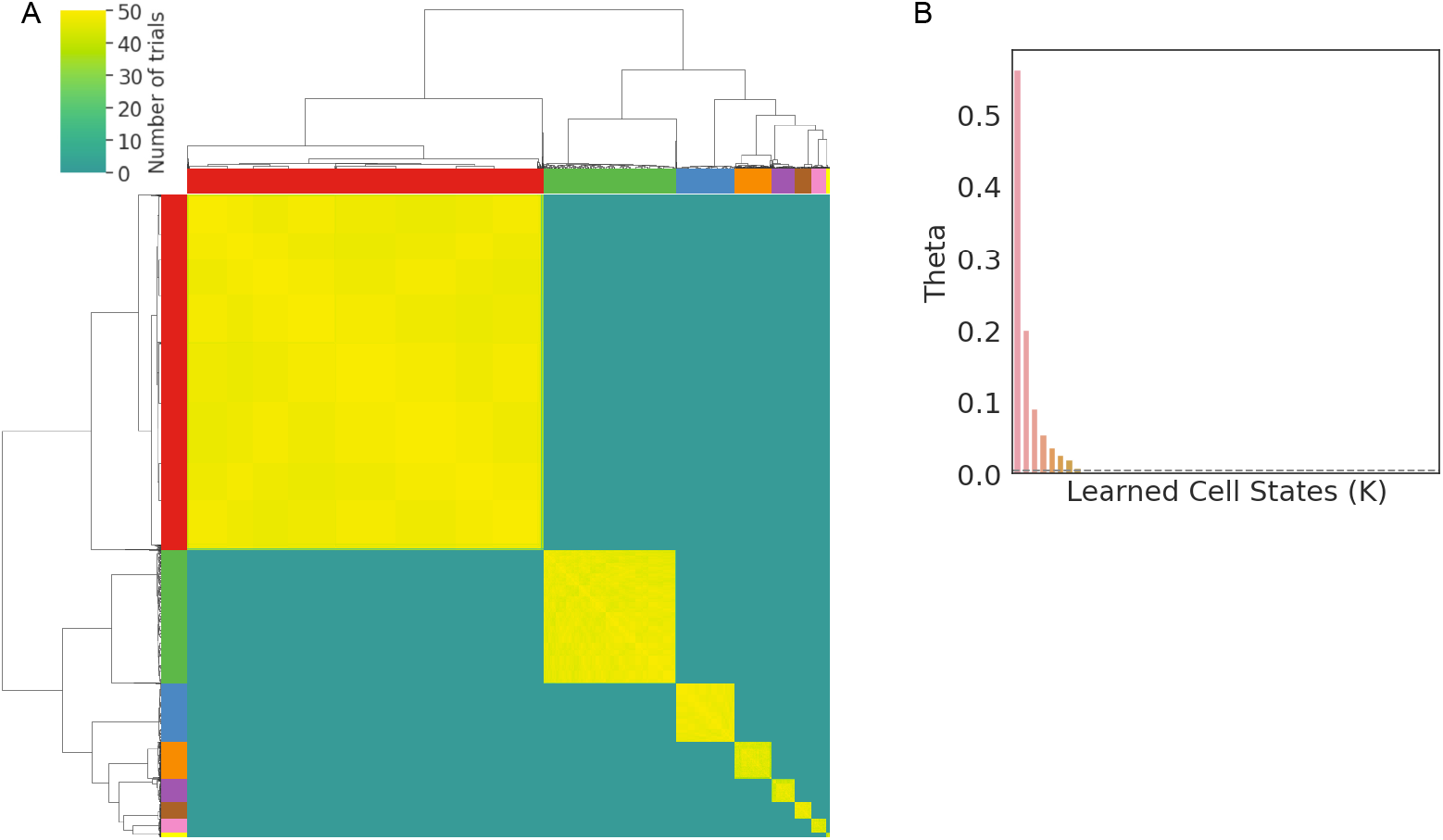
(A) Hierarchical clustering of cell-cell co-assignments across 50 independent trials on simulated data. Color bars indicate pre-assigned cell types. (B) Learned values for *θ* across 50 cell states. Dashed grey line indicates 0.05%.

## 7 Application to Tabula Muris Brain data

### 7.1 Model consistency in cell assignments and association with known cell types

Smart-seq2 data for individual cells across eight brain cell types in Tabula Muris was processed as described in **Section 2**. Inton clusters were then annotated with M10-PLUS genes. In this reference-aware mode, we detected 16,191 junctions across 5,433 intron clusters and 4,135 genes.

We initially assessed Leaflet’s clustering consistency across ten random initializations by calculating how often each pair of cells were assigned to the same cell state (to account for label switching between runs). We set *K* to 10 for this assessment. The results revealed a strong tendency for the same cell type pairs to be co-assigned, highlighting clear cluster structure in the data (Adjusted Rand Score = 0.80, **Figure 4**). We then selected the trial with the highest ELBO from the ten runs. Inspecting the posterior mean *thêta* = *γ/* Σ_*k*_ *γ*_*k*_, we found that all *K* (10) cell states appeared in at least 5% of cells (**Figure 8A,B**). Notably, most cell states predominantly corresponded to one or two closely related cell types (**Figure 5A,B**). For example, Cell States 1 and 5 were predominantly enriched in macrophages and microglia (**Figure 5D**). These results suggest that Leaflet is able to recover biologically meaningful distinct cell-types using only splicing patterns.

**Figure 4:**
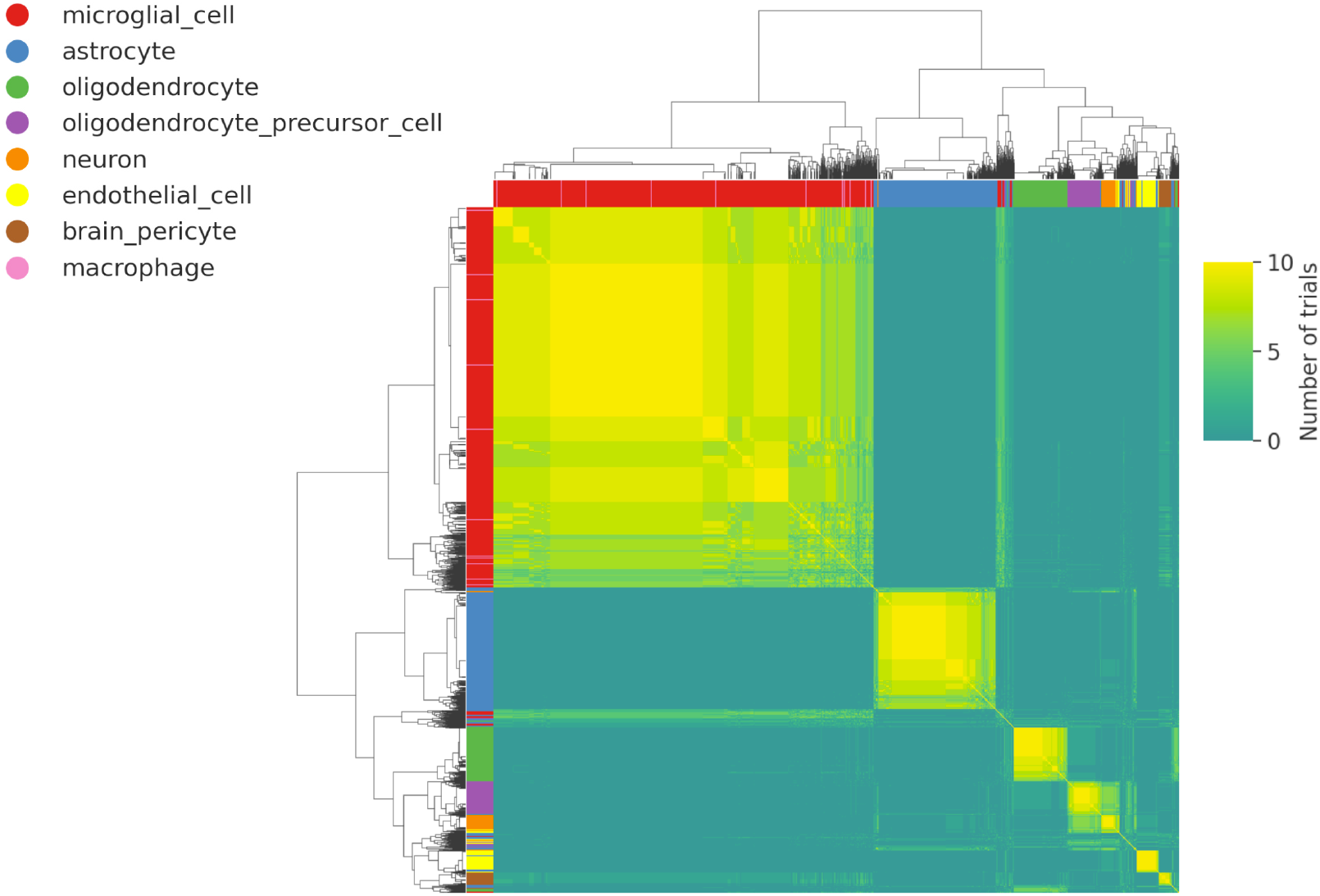
Hierarchical clustering of cell pair co-assignments across ten independent trials (n=5,200 randomly sampled cells are shown).

**Figure 5:**
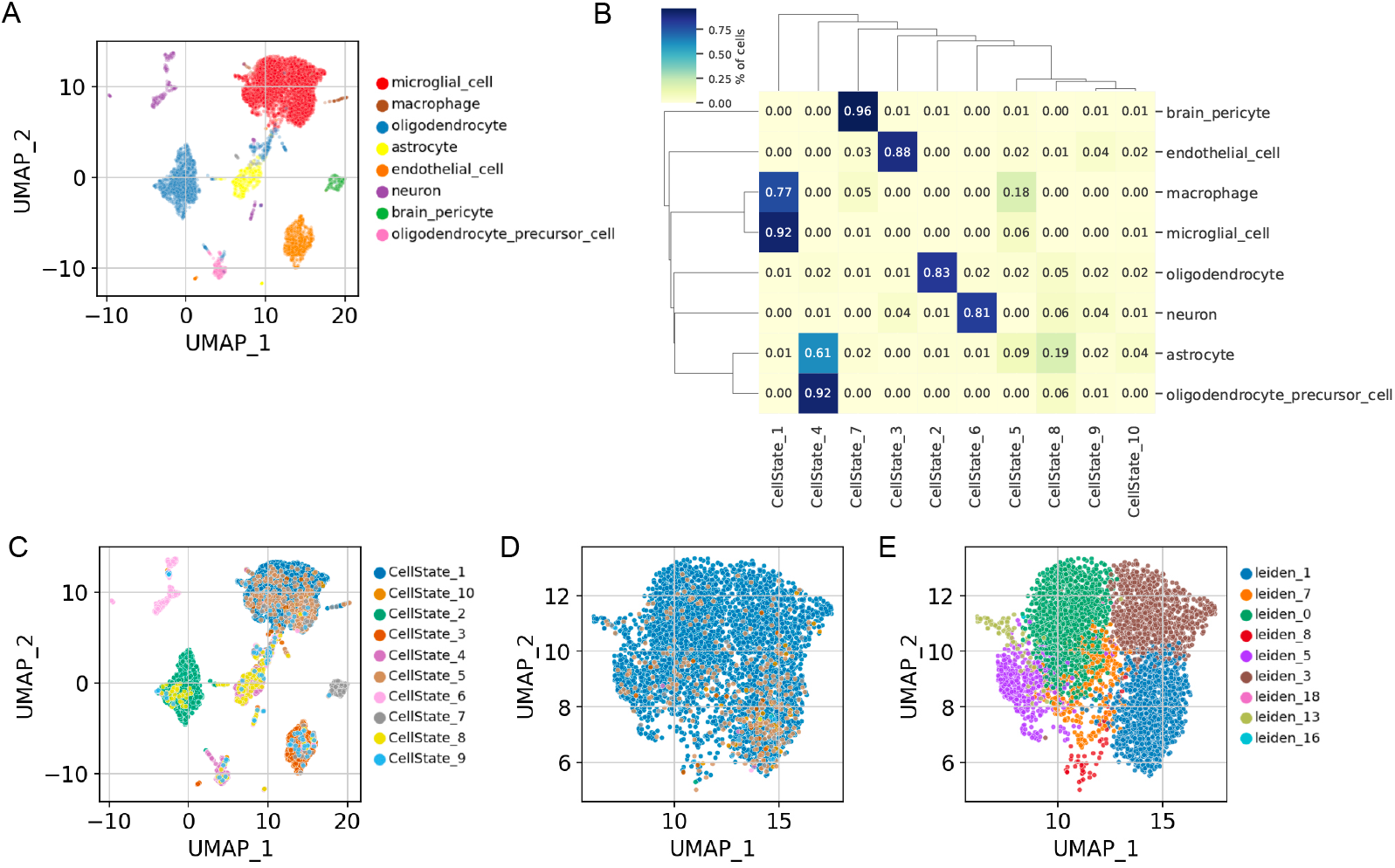
(A) UMAP based on PCA analysis of gene expression overlaid with known cell types. (B) Percentages of cell types belonging to each Leaflet cell state.(C) Leaflet cell state assignments overlaid on UMAP from (A). (D) Subset of UMAP plot is shown for predefined microglial cells (same coour legend as C). Cells are coloured by Leaflet cells states where the first and fifth state make up the majority of this population. (E) Same cells as in panel D but coloured by Leiden clusters defined by total gene expression.

### 7.2 Cell state specific splicing events shape cellular diversity

Cellular heterogeneity may be shaped by both gene expression and AS patterns, the latter of which is less well understood. We first evaluated the learned *J* × *K* matrix of Leaflet *ψ*s as described above (**Section 6.2**). We compared these learned *ψ* values to empirical (observed) junction usage ratios (per cell), both of which exhibited peaks at 0 and 1 (**Figure 6A,B**).

**Figure 6:**
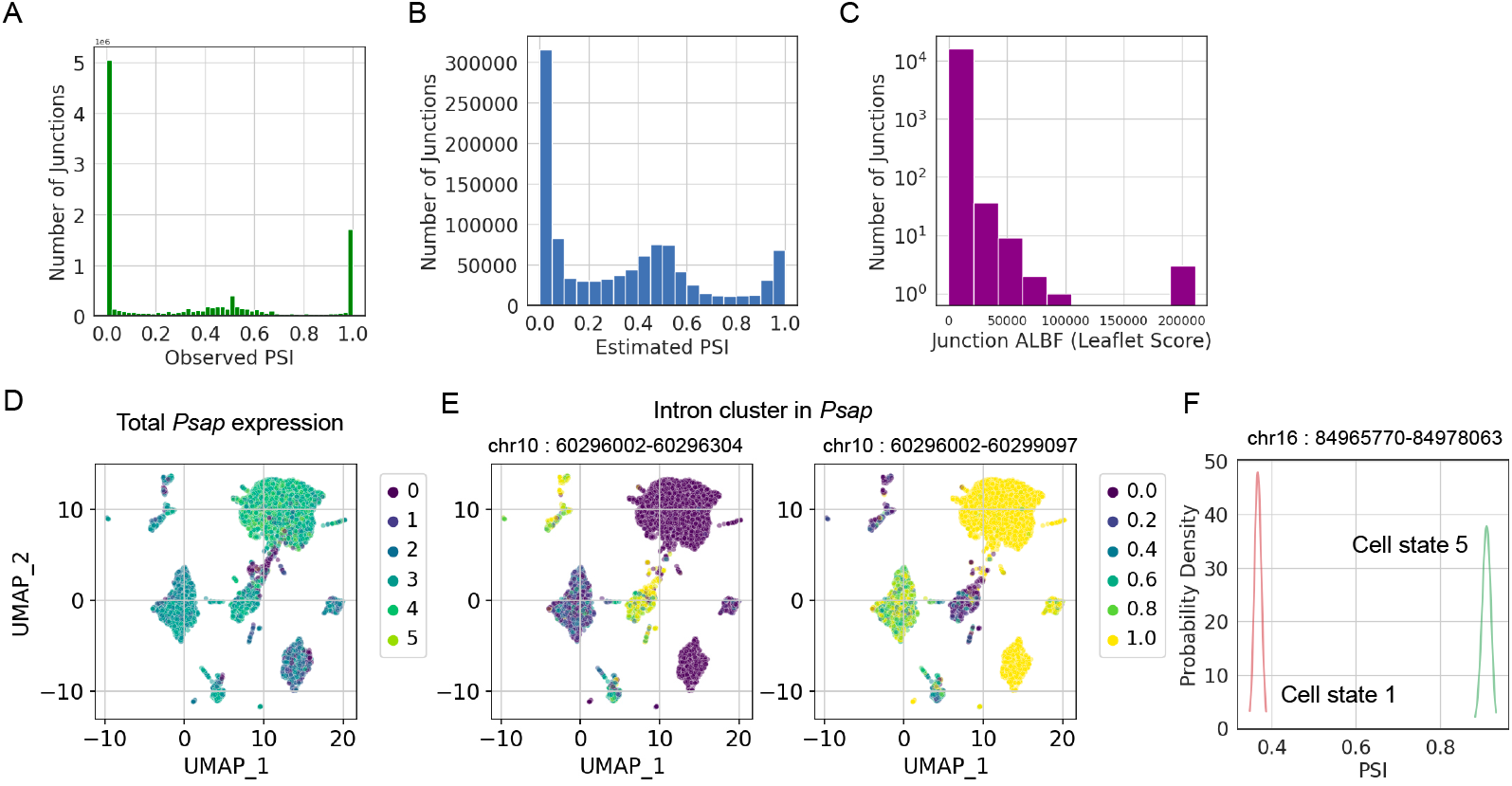
(A) PSI values for observed junction-cell pairs (*y*_*cj*_*/T*_*cj*_). (B) Distribution of Leaflet 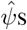s for junction-cell state pairs. (C) Junction ALBF scores across all cell states. PSI_*jk*_ are shown only for those junctions and cell states in which *α*_*jk*_ + *β*_*jk*_ ≥ 10 to exclude those with very high uncertainty (B,C).(D) *Psap* normalized gene expression values across cell types suggests it is not a marker gene. (E) An example of an intron cluster in the gene *Psap*, consisting of two junctions with high ALBF scores, showing variable AS across cell types. (F) Posterior distributions Beta(*α*_*kj*_, *β*_*kj*_) are shown for a junction mapping to the gene *App* in cell states 1 and 5.

We then obtained Leaflet ALBF scores for each junction, where higher scores indicate a junction is more variable across all *K* cell states (**Figure 6C**). We found that of the top 10% scoring junctions, less than 7% mapped to any marker genes, suggesting this strong splicing variation is mostly independent of expression. The top two scoring junctions were in the same intron cluster (corresponding to an alt 3’ splice site) in the *Psap* gene. One splice site is used predominately in pericytes and oligodendrocytes, while the other is used in neurons and astrocytes (and interestingly oligodendrocyte precursors show a mix of both, **Figure6D**,**E**).

We next calculated Leaflet ALBF scores across only two cell states of interest (cell states 1 and 5), corresponding to microglial cells. One of the top scoring junctions mapped to the gene *App* and was highly used in cell state 5 compared to cell state 1 (**Figure 6F**). This highlights that there may be a significant level of AS heterogeneity even within a well defined cell type such as microglia. We note that these cells are not directly linked to total gene expression based subpopulation of cells as determined by Leiden clustering (**Figure5**E) and may correspond to complementary levels of information.

## 8 Discussion

In this work, we introduce Leaflet, a novel, flexible approach to identify AS-driven cell states, a dimension of cellular diversity that is often unexplored. By focusing on local splicing variation at the single-cell level and comparing the differential splicing observed across learned cell states with those observed in pre-defined cell types, Leaflet reveals AS heterogeneity even within established cell types such as microglia. Our preliminary findings suggest that the landscape of cell-type specific differential splicing is not only a proxy for differential gene expression, but an additional, orthogonal layer of regulatory information.

While the current application to brain cells sequenced with Smart-seq2 has proven valuable, we plan to expand our method to analyze high-throughput datasets from techniques such as SPLiT-seq [16] and EasySci [17]. Although long-read scRNA-seq might ultimately be the definitive approach for deciphering isoform variation, short-read protocols look likely to remain dominant in the short to medium term.

While any biological insights made herein require additional validation in external datasets, we present a new approach and highlight several examples for further study. Leaflet is fast to run and requires minimal input from users.

## 9 Acknowledgements

This material is based upon work supported by the National Science Foundation under Grant No. DBI2146398. Any opinions, findings, and conclusions or recommendations expressed in this material are those of the authors and do not necessarily reflect the views of the National Science Foundation.

## 10 Appendix

### 10.1 Evidence Lower Bound (ELBO)

The ELBO can be decomposed as

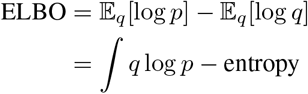

where the former term for Leaflet’s mixture model is

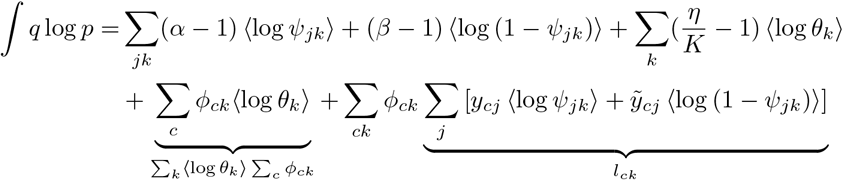

where *ϕ*_*ck*_ = ⟨*z*_*c*_ = *k*⟩ and 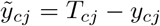.

### 10.2 Splicing variation along a simulated continuous trajectory

Here we compare Leaflet and Psix, utilizing data that was simulated to align with continuous cell state trajectories. The simulated dataset consists of 5,000 genes and 1,000 cells that follow a one-dimensional trajectory where 40% of genes have variable splicing. Each gene was simulated to have two isoforms, one with and without a cassette exon. For exons marked as “non-cell state associated”, their PSIs were simulated to be the same across all cells (**Figure 7**). This dataset corresponds to the simulated data used in the Psix manuscript [11]. The Kruskal-Wallis test was used as a baseline. Given the junction-centric approach of Leaflet, gene-level Leaflet ALBF scores were derived by summing the ALBF scores for relevant junctions. Importantly, Leaflet was not designed for continuous, trajectory-like cell states yet it obtained accurate estimates even on such data (**Table 3**). Additionally, Psix runtime was 22.2 minutes while Leaflet inference took only 35 seconds with 100 CAVI iterations (all methods tested on CPU).

### 10.3 Low dimensional representation of Tabula Muris brain cells using gene expression

We utilized total gene counts obtained with FeatureCounts [18] for these cells to obtain gene expression based low dimensional representations for downstream analysis. We used the Scanpy pipeline with recommended settings for quality control of cells and genes [19] and total count based normalization. Principal Component Analysis (PCA) was run on 7,560 cells and 3,837 genes. Uniform Manifold Approximation and Projection (UMAP) was run on the first 40 principal components (PCs) to obtain a low dimensional transcriptomic representation of the cells (**Figure 5A**). Marker genes (n=30) were obtained for each of the nine cell types in the data using the built in T-test approach.

### 10.4 Supplementary Figures

**Figure 7:**
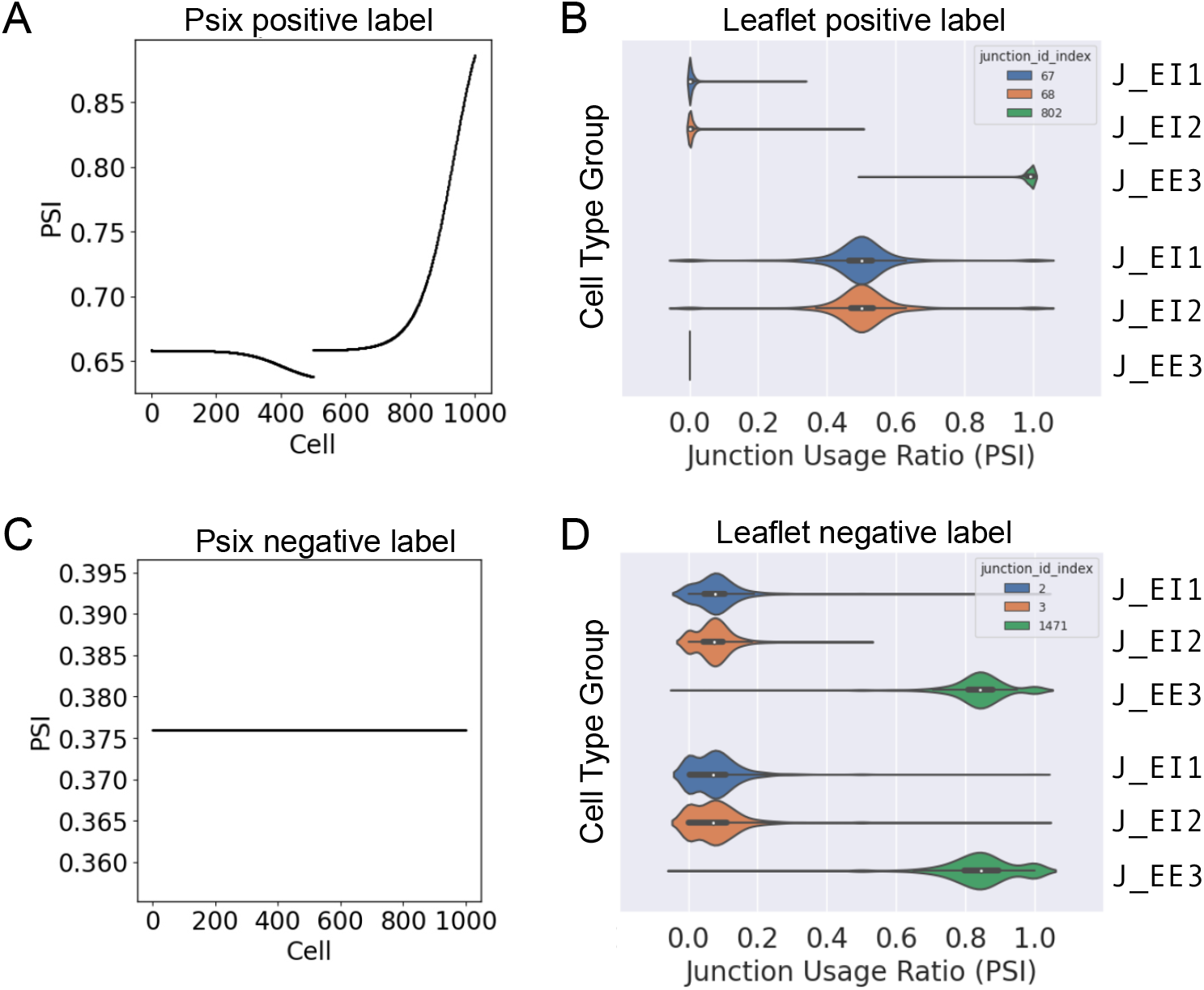
Simulated PSIs from Psix (left) and Leaflet (right). Simulated differential splicing is shown in the top panels for positively labelled events. Psix simulates cells along a trajectory. A cell state associated exon is shown in **A** and a cell type associated intron cluster composed of three junctions is shown in **B**. Bottom panels show instances of simulated PSI values for non-differentially spliced exons (**C**) or intron clusters (**D**). *J*_*EI*_ : Exon inclusion junction. *J*_*EE*_: Exon exclusion junction.

**Figure 8:**
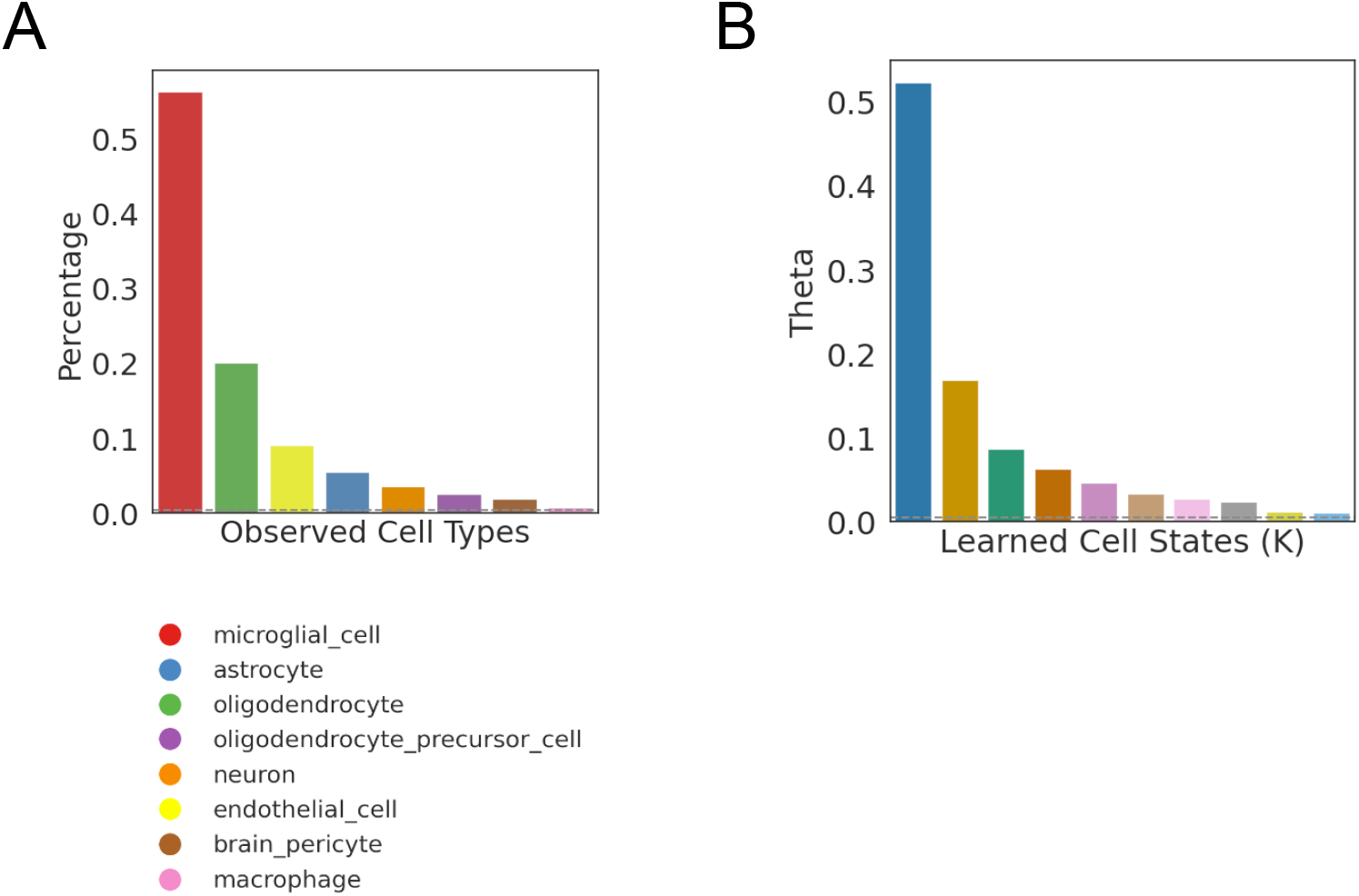
(A) Distribution of predefined cell types in the dataset. (B) Distribution of *θ* learned for each cell state.

## Notes

### Competing Interest Statement

The authors have declared no competing interest.

### Summary of Updates

An acknowledgements section was added to reflect funding sources.

